# The tumor suppressor protein PTEN undergoes amyloid-like aggregation in tumor cells

**DOI:** 10.1101/2020.11.30.402115

**Authors:** Filip Claes, Elisabeth Maritschnegg, Greet De Baets, Aleksandra Siekierska, Mirian Saiz Rubio, Meine Ramakers, Emiel Michiels, Frederik De Smet, Jeroen Depreeuw, Ignace Vergote, Adriaan Vanderstichele, Annick Van den Broeck, Siel Olbrecht, Els Hermans, Frédéric Amant, Diether Lambrechts, K. Peter R. Nilsson, Frederic Rousseau, Joost Schymkowitz

**Author notes:** Corresponding authors: Frederic Rousseau and Joost Schymkowitz.

## Abstract

Protein aggregation is an underappreciated mechanism that may contribute to the loss- and oncogenic-gain-of-function of mutant tumor suppressors such as p53 and axin. In the present study, we describe amyloid-like aggregation behaviour of the second most frequently mutated tumor suppressor in human cancer, PTEN. *In silico* analysis revealed a particularly high aggregation vulnerability for this protein, which was corroborated by *in vitro* aggregation assays. In cultured tumor cells, we found that under stress conditions, PTEN readily undergoes amyloid-like aggregation as a result of mutation. However, we also show that severe dysregulation of protein homeostasis may lead to aggregation of wild-type PTEN. These observations were supported by a small survey of patient-derived uterine tumor tissues, which found that more than 25% of tumors analyzed displayed wild-type PTEN aggregation. Finally, in an exploratory clinical study we found that PTEN aggregation status was correlated with a decline in clinical outcome. Our findings establish that the tumor suppressor PTEN is highly aggregation-prone and our work suggests that protein aggregation might be an underestimated but prevalent component of cancer cell biology.

## Introduction

Phosphatase and tensin homolog or PTEN is the second most commonly mutated tumor suppressor in human cancers (Cetintas and Batada, 2020). Its tumor suppressive role is largely dependent on its ability to downregulate the growth and survival promoting phosphatidylinositol-3-kinase (PI3K)/AKT pathway (Hoxhaj and Manning, 2020). Here, PTEN drives the dephosphorylation of the AKT-activating substrate PIP3 to PIP2 through its lipid phosphatase activity (Maehama and Dixon, 1998). Nevertheless, alternative functions have been shown to contribute significantly to PTEN’s tumor suppressive function, including control of cellular metabolism and maintenance of genome stability (reviewed in (Lee et al., 2018)).

PTEN function is deregulated in a large proportion of human cancers through different mechanisms, including somatic mutations, gene silencing, or epigenetic mechanisms (Cetintas and Batada, 2020). Although a clear hotspot mutation site is observed in the phosphatase domain, cancer-associated mutations may occur in all PTEN domains, supporting the observation that different PTEN functions, associated with different protein domains, contribute to the tumor suppressive role of PTEN (https://cancer.sanger.ac.uk/cosmic/gene/analysis?ln=PTEN#gene-view). Interestingly, tumor syndromes associated to germline mutations in *PTEN* in man and mouse (Eng, 2003; Mester et al., 2011) suggest that expression of a mutant with reduced activity may result in a worse prognosis than when PTEN expression is simply lost, suggesting PTEN might also deploy oncogenic gain-of-function under certain conditions (Muller and Vousden, 2013) (Papa et al., 2014; Spinelli et al., 2015; Wang et al., 2010).

Protein aggregation is typically associated to neurodegenerative and amyloidogenic disorders, where the abnormal accumulation, aggregation and inclusion body/extracellular deposit formation of disease-specific proteins are defining hallmarks of the disease (Lim, 2019; Soto and Pritzkow, 2018). However, protein aggregation might also contribute to cancer development and progression. Indeed, we and others have shown that aggregation of mutant p53 tumor suppressor can contribute to cancer by inducing dominant loss-of-function of wild-type tumor-suppressing activity, but also gain-of-function by co-aggregating with various cellular components, including other tumor suppressors and cell cycle regulators (Ano Bom et al., 2012; De Smet et al., 2017; Forget et al., 2013; Joerger and Fersht, 2008; Lasagna-Reeves et al., 2013; Levy et al., 2011; Wang and Fersht, 2012; Wilcken et al., 2012; Xu et al., 2011; Yang-Hartwich et al., 2015). Furthermore, point mutations in the scaffold protein Axin were shown to promote its aggregation resulting in an oncogenic gain-of-function through rewiring of the Wnt signaling pathway (Anvarian et al., 2016). Protein aggregation could therefore constitute a novel unexplored mechanism that is exploited by tumor cells to drive their progression.

Recently, an *in silico* survey of the wild-type PTEN protein sequence and 1523 of its clinically relevant mutant sequences was performed to assess changes in intrinsic aggregation propensity of the mutant polypeptide sequences. This analysis revealed that while the PTEN wild-type protein already shows intrinsic aggregation propensity, 18% of the disease-associated PTEN mutants further increased this propensity significantly (Palumbo et al., 2020). Here, we further corroborate these theoretical findings and show experimentally that both wild-type and mutant PTEN can indeed form oligomeric and amyloid-like aggregates in cultured tumor cells *in vitro.* Furthermore, in a set of patient-derived uterine tumor samples, we found that more than 25% of tumors analyzed displayed PTEN aggregation. However, PTEN aggregation in this set appeared to be almost exclusively associated to tumors that were wild-type for PTEN. Finally, in an exploratory clinical study we found that PTEN aggregation status was correlated with a decline in clinical outcome.

Our findings establish experimentally that the tumor suppressor PTEN is indeed highly aggregation-prone and that wild-type PTEN protein aggregation might constitute a novel mechanism of PTEN inactivation during tumor development or progression.

## Results

### PTEN is an aggregation prone tumor suppressor protein

Protein aggregation is the self-association of a protein through non-native interactions, which are usually mediated by short hydrophobic stretches in the primary protein sequence, termed aggregation prone regions (APRs) (Rousseau et al., 2006). Previously, it was shown that mutations, which are predicted to increase the aggregation propensity of proteins are enriched in cancer (De Baets et al., 2015), suggesting that protein aggregation plays a greater role in tumorigenesis than expected. As aggregation is expected to disrupt normal function of the target protein, we further narrowed down this analysis to a non-exhaustive list of commonly mutated tumour suppressors, whose normal function is to protect the cell from carcinogenesis. To this end, we first performed an *in silico* analysis of the intrinsic aggregation propensity of their wild-type protein sequences using TANGO (Fernandez-Escamilla et al., 2004) (Table 1). Our analysis showed that many of the tumor suppressors we analysed carry particularly strong APRs (score >60), which is consistent with previous findings (De Baets et al., 2015). From this list we next selected PTEN for further in-depth characterization as (i) PTEN is considered to be the second most commonly mutated gene in sporadic cancers after p53, and (ii) the PTEN protein was previously shown to be only marginally stable and might therefore be particularly susceptible to aggregation (Johnston and Raines, 2015), in line with predictions from others (Palumbo et al., 2020). Detailed TANGO analysis of PTEN showed that there are four APRs with a TANGO score >60 (Figure 1a). Two of them localize to the phosphatase domain, while the remaining two are localized in the C2 domain (Figure 1b). To assess the aggregation propensity of the PTEN protein, we recombinantly expressed and purified the N-terminal 274 amino acids of PTEN (N-PTEN) (Ramaswamy et al., 1999) containing the complete phosphatase domain to which the majority of the most destabilizing PTEN mutations localize. N-PTEN readily formed amorphous aggregates as evidenced by transmission electron microscopy (Figure 1c)

**Figure 1.**
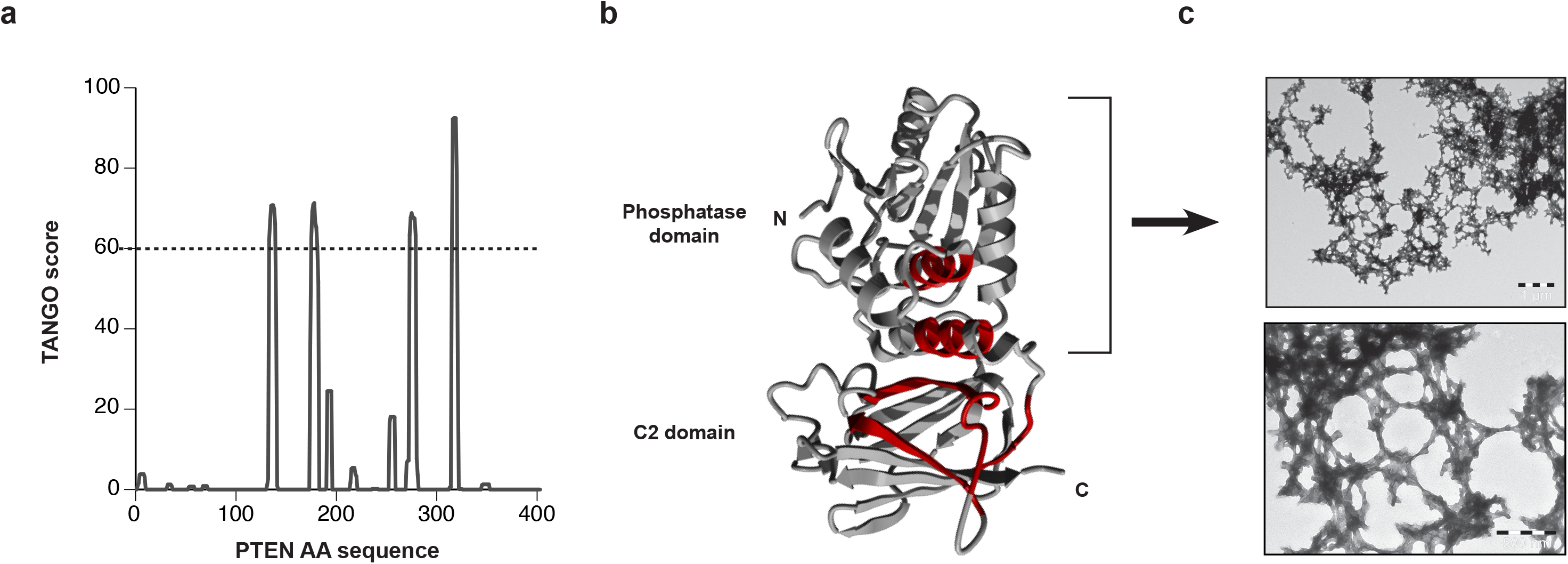
*In silico* and *in vitro* analysis of PTEN aggregation propensity. **a.** Plot of the TANGO scores over the PTEN amino acid sequence, predicting regions that are prone to aggregation (aggregation prone regions, APRs). Four different regions display a TANGO score >60 suggesting that these regions have strong tendency to aggregate. **b.** Overview of the PTEN protein structure in which the APRs where highlighted in red. Two of the APRs are located in the β-strand-rich C-terminal C2 domain (bottom), while the other two are located in the N-terminal phosphatase domain (top). **c.** Transmission Electron Microscopy (TEM) images of 5 DM N-PTEN in 4M urea in PBS after 24 h incubation at room temperature and negatively stained with 2% (w/v) uranyl acetate.

**Table 1.** Bioinformatic analysis of the aggregation propensity of tumor suppressor proteins

| uniprot | tumor suppressor | number of APR with TANGO>60 | max scoring APR |
| --- | --- | --- | --- |
| RB_HUMAN | Retinoblastoma-associated protein | 8 | 99 |
| TSC2_HUMAN | Tuberin | 13 | 98 |
| MLH1_HUMAN | DNA mismatch repair protein Mlh1 | 2 | 97 |
| CADH1_HUMAN | Cadherin-1 | 10 | 96 |
| PTC1_HUMAN | Protein patched homolog 1 | 20 | 96 |
| BRCA2_HUMAN | Breast cancer type 2 susceptibility protein | 6 | 95 |
| MSH2_HUMAN | DNA mismatch repair protein Msh2 | 3 | 95 |
| NF1_HUMAN | Neurofibromin | 18 | 95 |
| TSC1_HUMAN | Hamartin | 2 | 91 |
| BRCA1_HUMAN | Breast cancer type 1 susceptibility protein | 3 | 90 |
| FBXW7_HUMAN | F-box/WD repeat-containing protein 7 | 2 | 89 |
| P53_HUMAN | Cellular tumor antigen p53 | 1 | 89 |
| DCC_HUMAN | Netrin receptor DCC | 4 | 86 |
| CD2A1_HUMAN | Cyclin-dependent kinase inhibitor 2A, isoforms 1/2/3 | 1 | 82 |
| PTEN_HUMAN | PTEN | 4 | 76 |
| APC_HUMAN | Adenomatous polyposis coli protein | 1 | 70 |
| MERL_HUMAN | Merlin | 2 | 70 |

### cancer-associated mutations affect the aggregation propensity of tumor suppressors

To assess whether cancer-associated inactivating PTEN mutations might be linked to aggregation, we analysed the predicted effect of a PTEN mutation set - as taken from the Humsavar db - on both protein stability (using the FoldX force field (Schymkowitz et al., 2005)) and intrinsic strength of APR sequences (using TANGO). We then mapped the aggregation propensity spectrum of these disease-associated PTEN mutations on a mass-plot, combining changes in intrinsic TANGO scores and changes in thermodynamic stability (Supplemental Figure 1a). As observed by others (Emily, 2019), several PTEN mutations are predicted to increase vulnerability of the protein to aggregation through either (i) destabilizing its native structure thereby increasing exposure of APRs, (ii) increasing the intrinsic self-association propensity of APRs, making them more prone to self-interaction when exposed or (iii) a combination of the two.

From this analysis we selected three missense mutations, all in the N-terminal phosphatase domain, for further analysis: (i) R47G, a control mutation with no significant effect on stability and aggregation propensity; (ii) R173P, a mutation predicted to moderately decrease stability and increase APR strength; and (iii) C136Y, a mutation predicted to severely decrease stability and increase APR strength. Like for other tumor suppressor genes – except p53 - nonsense and frame-shift mutations are common mutation types for PTEN (Supplemental Figure 1b). Indeed, nonsense mutations account for 16% and frame-shift mutations for 28% of all cancer-associated PTEN mutations (data taken from Cosmic db). These types of mutations may result in translation of destabilized truncated PTEN proteins in case of a premature stop (potentially increasing exposure of endogenous APRs) or even of abnormal de novo protein products in case of frame-shifted translation (potentially creating novel APRs). Therefore, we also included some of the most common mutations from each category, i.e. R233X and 800delA, respectively, in our study.

### PTEN mutant stability in physiological conditions

To assess whether the selected mutants alter the subcellular distribution of the PTEN protein, we transiently expressed them in the HeLa cervical adenocarcinoma cell line. Qualitative analysis of the immunofluorescent staining pattern showed a diffuse cytosolic distribution for all missense mutants, similar to wild-type (Figure 2a and Supplemental Figure 2a). Although similar observations were made for the truncating frameshift and nonsense mutants on the subcellular distribution of the protein, the number of cells staining positive was clearly lower. This observation was corroborated by high-content analysis, which revealed that while PTEN wild-type and all mutant proteins were expressed at similar levels in expressing cells (Supplemental Figure 2b), the fraction of cells staining positive for the R233X nonsense and the 800delA frameshift mutants were detected >5 times less frequently as compared to the missense mutants, indicating that these mutants were effectively cleared by most cells (Supplemental Figure 2c).

**Figure 2.**
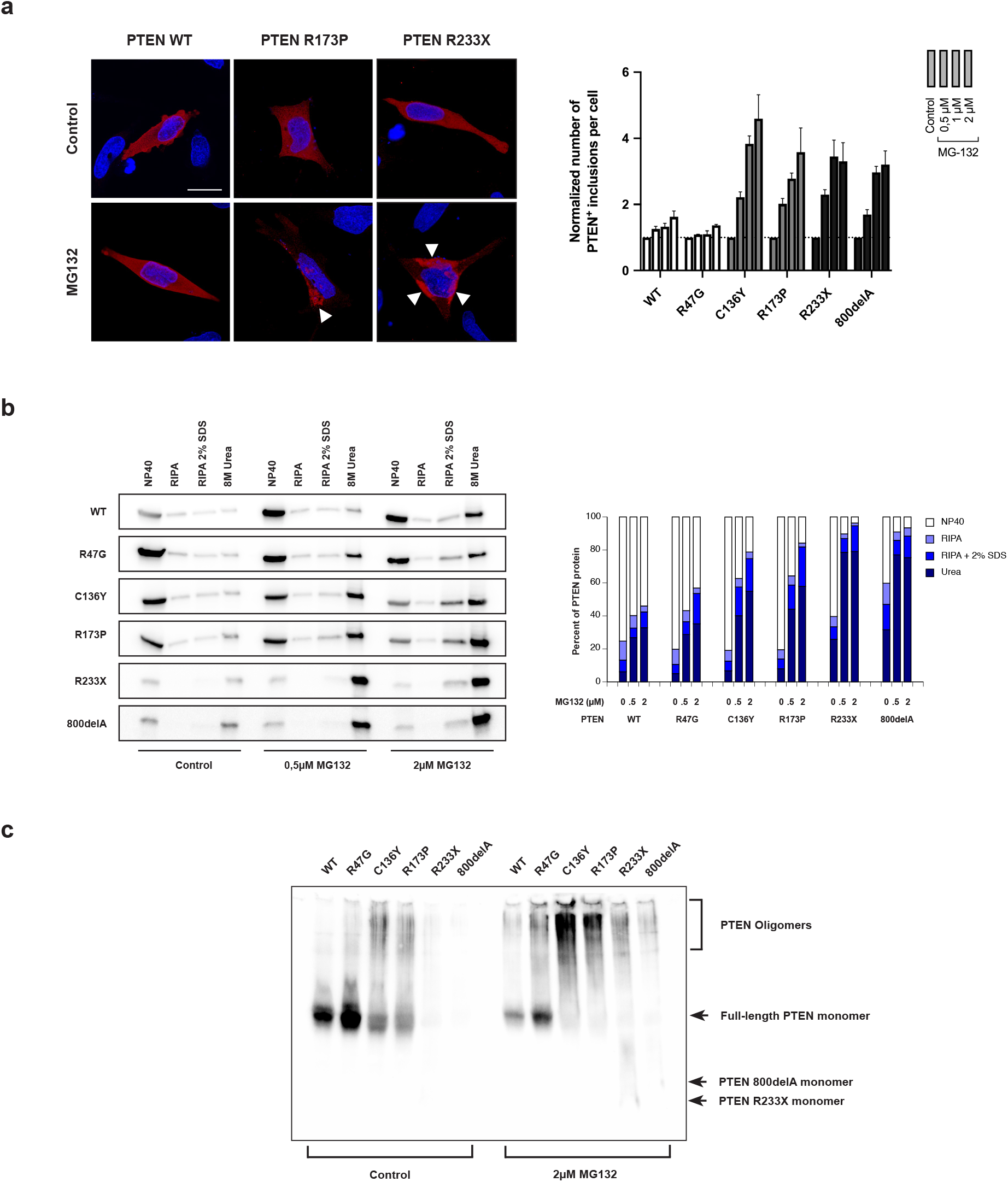
PTEN mutants can form aggregate-like structures upon proteasomal inhibition. **a.** (left panels) Immunofluorescence of PTEN WT and the R173P and R233X mutants in HeLa cells under optimal cell culture conditions (top row) and after proteasomal inhibition with MG132 treatment (bottom row). Under control conditions PTEN shows a diffuse distribution throughout the cell. However, upon MG132 treatment, the staining of the PTEN mutants R173P and R233X becomes localized to protein inclusions, while most of the PTEN WT protein retains its diffuse staining pattern. Scale bar represents 20μm. (right panel) High-content analysis of protein inclusion formation of PTEN WT and the selected mutants in HeLa cells under control conditions and proteasomal inhibition (MG132). Treatment of the destabilizing mutants with increasing doses of MG132 resulted in a steep, dose-dependent increase in the average number of protein inclusions per cell. For PTEN WT and the control mutant R47G, however, only a very mild increase in the number of inclusions. **b.** Assessment of PTEN protein solubility using the serial extraction assay. (left panel) Western blots showing the distribution of the WT and mutant PTEN proteins over the different solubility bins under control and MG132-treated conditions. (right panel) Protein content in each solubility bin was quantified densitometrically and expressed as percentage of total protein for each condition. Treatment of transfected HeLa cells with MG132 resulted in a dose-dependent decrease in protein solubility as evidenced by a shift of the most soluble NP40 fraction (white bar fragments) to the most insoluble urea fraction (dark bleu bar fragments). Consistent with inclusion formation, this shift was most prominent for the destabilizing and truncation mutants. **c.** Non-reducing and less-denaturing SDS PAGE and Western blot for PTEN WT and mutants in the soluble NP40 fraction. Under control conditions a high-molecular weight smear - indicative for oligomeric species - was observed for the destabilizing C136Y and R173P PTEN mutants but not for WT and the stable R47G mutant. The nonsense and frameshift mutants, 800delA and R233X respectively, were not reliably detected under these conditions. Upon treatment with MG132 all PTEN variants showed an increase in high-molecular weight smearing, showing the induction of oligomer formation upon proteasomal inhibition.

To assess gross differences in solubility indicative of aggregation between PTEN variants, we next performed a serial protein extraction protocol. Indeed, a defining hallmark of protein aggregates - when compared to native proteins - is their increased resistance to chemical denaturants and detergents. To assess this, we performed a serial extraction assay: proteins form PTEN-expressing HeLa cells were extracted in buffers of increasing denaturing strength, whereby the insoluble fraction in a given buffer is collected by centrifugation and re-suspended in the next buffer of the series. Aggregated proteins are then expected to only be solubilized in the more stringent buffers. For both wild-type PTEN and the missense mutants most of the protein (>70%) was detected in the most soluble fractions 24hrs after transfection. Consistent with the immunofluorescence, the R233X nonsense and the 800delA frame-shift mutant protein levels detected on Western blot were much lower as compared to the full-length mutants, however, they appeared to be less soluble, with most of the protein now distributed to the more insoluble fractions (Figure 2b).

Finally, to assess the presence of soluble PTEN oligomers in the NP40 fraction, we used a non-reducing, mildly denaturing SDS PAGE approach (Figure 2c). This analysis showed that under control conditions PTEN WT and R47G was resolved at the expected molecular weight, and thus formed no or little oligomeric species. However, for PTEN C136Y and R173P an additional high molecular weight smear was observed, indicating the presence of soluble oligomers. The nonsense and frameshift mutants, R233X and 800delA, respectively, were not detected under these conditions.

### Proteasomal malfunction triggers PTEN amyloid-like aggregation

Cancer cells tailor the proteostatic machinery to their needs, supporting uncontrolled survival and growth. Unusually high proteasomal activity, for example, appears to be a common feature in cancer cells and a prerequisite for their survival (REF). However, as aging, which is associated to a functional decline in the proteostasic machinery, is one of the biggest risk factors for developing cancer (Ershler and Longo, 1997; Falandry et al., 2014), we set out to study the behaviour of the PTEN variants in conditions of decreased proteostatic fitness. To this end, we first explored whether decreased proteosomal function modifies the aggregation properties of the PTEN variants. Immunofluorescent analysis showed that PTEN staining became increasingly confined to inclusions after exposure to increasing doses of the proteasomal inhibitor MG132 (Figure 2a and Supplemental Figure 2a). High-content quantitative analysis showed that while this effect was limited for wild-type PTEN and the control mutant R47G, all other mutants showed a steep increase in the percentage of cells showing punctuate PTEN staining with increasing MG132 dose.

To study whether the observed inclusions correlate with aggregation-specific parameters, we first performed the serial extraction assay, as described above. This assay showed that treatment of PTEN transfected cells with MG132 caused a dosedependent decrease in solubility for all constructs tested. However, paralleling the formation of protein inclusion-like structures, the decrease in solubility was more steep for the destabilizing mutants as compared to wild-type PTEN and the control mutant R47G (Figure 2b). Proteasomal inhibition also increased the level of oligomeric species in the NP40 fraction, as evidenced by a clear increase in high-molecular weight smears as compared to the control condition for all PTEN variants tested (Figure 2c), when assessed by reduced-denaturing SDS-PAGE.

To further characterize the nature of the insoluble PTEN protein fraction, we first assessed the particle size of the SDS-resistant PTEN protein fraction. To this end, we performed a filter trap assay. Specifically, PTEN transfected HeLa cells were lysed in a high SDS (1%) extraction buffer and lysates were filtered through a 0,22 μm filter. Proteins retained on the filter were recovered by stringent washes using boiling in 2% SDS followed by an 8M urea extraction step. PTEN levels in the flow-through and trapped fractions were quantified using Western Blot. This analysis showed that MG132 treatment induced the formation of SDS-resistant high molecular weight PTEN particles that appear to be larger than 0,22μM. Although a significant increase was observed for wild-type and all tested mutant forms of PTEN, the respons was most prominent for the most destabilizing and/or aggregation prone mutants (Figure 3a). Notably, the effect appeared to be underestimated for the nonsense mutant R233X and the frameshift mutant 800delA. Indeed, to control for the recovery of the protein material from the filter, input levels were compared to flow-through and trapped levels. This showed that for both the R233X and 800delA mutants recovery of the trapped fraction appeared to be incomplete, indicating that these mutants were able to form very stable high-molecular weight species upon proteasomal inhibition.

**Figure 3.**
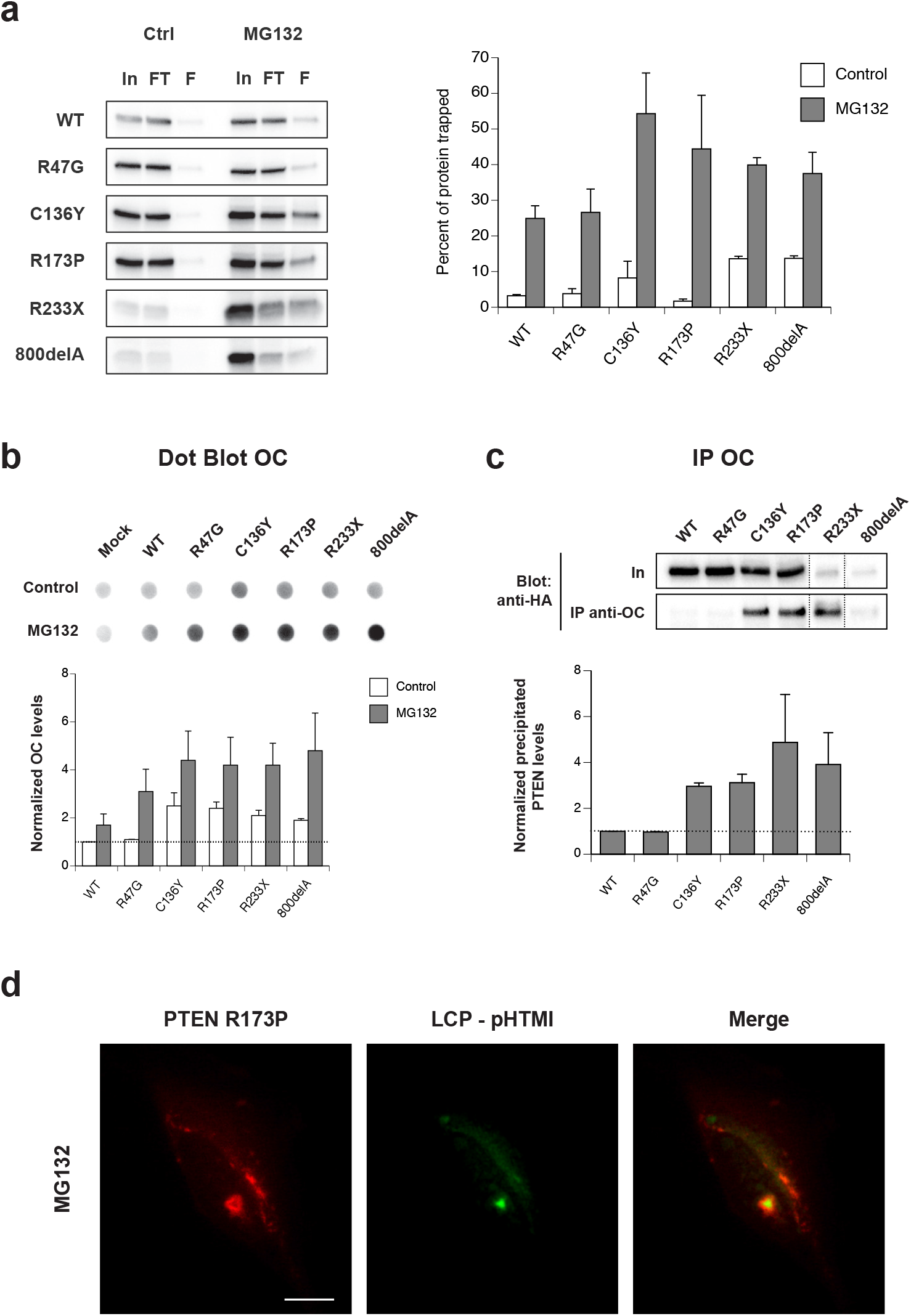
Characterization of PTEN aggregates formed upon proteasomal inhibition. **a.** Assessment of PTEN aggregate size by the filter trap assay. Cell lysates from PTEN transfected cells treated with or without MG132 were filtered through 0,22 μm membranes and input, flow-through and trapped fractions were blotted and probed for PTEN (left panel; In: Input, FT: flow-through fraction, F: fraction trapped on filter). Quantification of these blots revealed that the percentage of trapped PTEN protein increased markedly upon treatment with MG132 and the extent of this increase was largest for the destabilizing and truncation mutants. **b.** The structure of PTEN aggregates was probed by dot blot analysis with the amyloid fibril-specific antibody OC. Quantification of these blots showed that MG132 treatment induced the formation of OC-reactive structures in the lysates of PTEN transfected cells. **c.** To assess directly whether PTEN aggregates had an amyloid-like structure, the OC-reactive fraction was immunoprecipitated from PTEN transfected cell lysates and probed for PTEN. Precipitated PTEN levels were normalized to input levels and expressed relative to PTEN WT. **d.** The amyloid-like structure of PTEN aggregates was further confirmed by staining with the LCP dye pHTMI. Upon treatment of PTEN R173P transfected cell with MG132, inclusions were formed that stained positive for pHTMI, suggestive for the presence of fibril-like structures in the PTEN-positive inclusion. Scale bar represents 10 μm.

To gain more insight into the structural features of the PTEN aggregates formed upon proteasomal inhibition, we next characterized these PTEN aggregate species with a set of amyloid-specific probes. First, we performed dot-blot analysis using the fibril-conformation specific antibody OC (Kayed et al., 2007) on lysates of cells transfected with the different PTEN expression constructs. Under unstressed conditions, transfection of the destabilized and truncated mutants only caused a mild increase in OC signal over wild-type PTEN or the control mutant R47G, consistent with our other findings (Figure 3b). However, upon treatment with the MG132, there was a large increase in OC reactivity for all constructs tested. As observed before, this effect was most pronounced with the destabilizing and truncated mutants.

To assure that the observed increase in OC reactivity was directly attributable to the formation of PTEN fibril-like structures, we performed immunoprecipitations with the OC antibody in the same setting and probed for PTEN reactivity in the precipitated fractions. In control conditions, no PTEN could be detected in the precipitated fractions. However, in the MG132-treated lysates PTEN was precipitated with the OC antibody (Figure 3c) and the level of pull-down was most prominent for the destabilizing mutants. Finally, to show that the observed inclusions upon immunofluorescent staining are indeed the aggregated PTEN species observed in the biochemical analysis, we stained cells with luminescent-conjugated olithiophenes (LCOs) probes. Therefore, we screened a panel of different LCOs (Cieslar-Pobuda et al., 2014; Magnusson et al., 2015) against reactivity for cells transfected with PTEN R173P and treated with MG132 (not shown). Clear positive staining was observed for p-HTMI (penta-Hydrogen-Thiophene-Methylated-Imidazole) (Figure 3d), indicating that the observed protein inclusions upon immunofluorescent staining indeed correlate directly to the amyloid-like PTEN species. Together, these results show that upon proteasomal decline, PTEN can form high molecular weight, insoluble, amyloid-like aggregates.

### Tumor microenvironment-related stresses induce PTEN aggregation

Cancer cells are exposed to a variety of stresss in the tumor microenvironment such as hypoxia and hypoglycemia. These stresses modify the proteostasic fitness of the cancer cells, resulting in the accumulation of misfolded protein species. Therefore, we next assessed whether an hypoxia mimetic and glucose deprivation affect aggregation behavior of the PTEN variants.

To study the impact of hypoxia, we exposed transfected cells to NiCl_2_, a hypoxia mimetic (Xu et al., 2010), and assessed PTEN aggregation by performing serial protein extractions (Figure 4a). Exposing transfected cells to 1mM of NiCl_2_ for 36hrs results in a severe reduction in solubility for all PTEN variants. However, as with proteasomal inhibition, the extent of the reduction was only moderate for wild-type PTEN and the control mutant R47G, but considerably larger for the destabilizing mutants.

**Figure 4.**
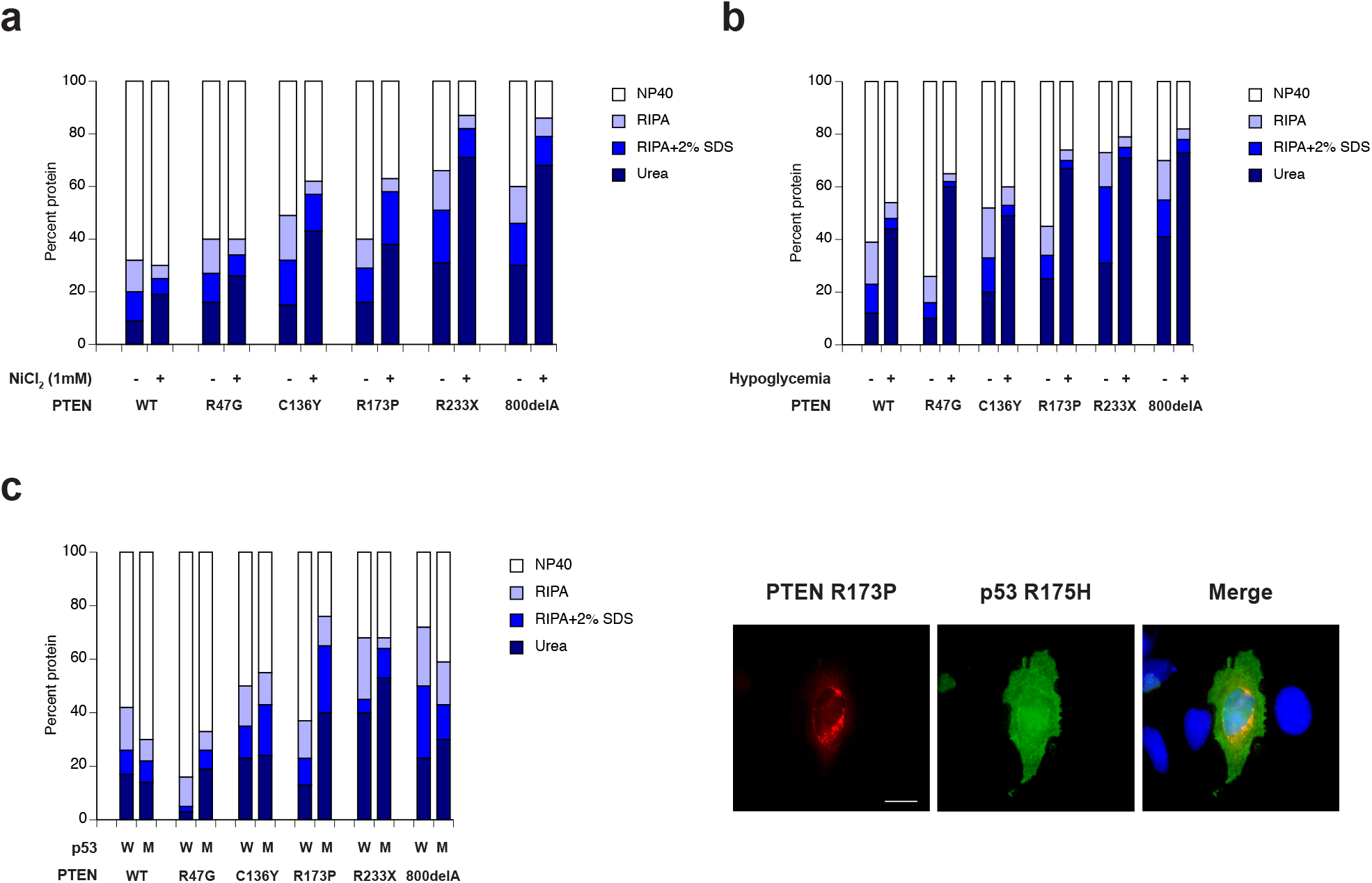
Exploring PTEN aggregation with other tumor-associated stresses. **a**. It is well established that the tumor microenvironment is short on oxygen and thus hypoxic. To mimic hypoxia, PTEN transfected cells were treated with 1mM NiCl_2_ for 48 hrs and aggregation was assessed by determining PTEN solubility in the serial extraction assay. This analysis showed that NiCl_2_ treatment caused a marked decrease in PTEN protein solubility for the destabilizing and truncation mutants, but not for PTEN WT and the control mutant R47G. **b.** High glucose consumption by tumor cells result in a hypoglycemic tumor environment. Culturing PTEN transfected cells in glucose-deficient medium for 48 hrs proved to be a strong trigger for PTEN aggregation as evidenced by the large shifts towards reduced PTEN protein solubility, even for PTEN WT and the control mutant R47G. **c.** Tumor cells may carry other aggregating proteins, as was shown for p53. To study PTEN behavior on an aggregation background, PTEN variants were co-transfected with p53 WT (W) or the aggregating mutant R175H (M) and PTEN protein solubility was assessed after 48 hrs of coexpression. This analysis showed that under these conditions only PTEN R173P showed a clear reduction in protein solubility when co-expressed with p53 R175H (left panel). Immunofluorescent staining showed that PTEN R173P became localized to inclusions upon co-expression with p53 R175H (right panels). These inclusions, however, did not stain positive for p53 indicating that there was no direct interaction between both. Scale bar represents 10μm.

Next, we assessed the effect of hypoglycemia on PTEN solubility, by minimizing the levels of glucose in the cell culture medium. Quantification of the different solubility fractions revealed that hypoglycemia is a remarkable powerful trigger for PTEN aggregation (Figure 4b). Indeed, in hypoglycemic conditions all PTEN variants tested, including wild-type and the R47G control mutant, showed a severe reduction in solubility as compared to when normal glucose levels are available. Given the potent effect of hypoglycemia on reducing PTEN protein solubility, we studied whether the reduction in solubility was paralleled by formation of protein inclusions as assessed by immunofluorescent staining (Supplemental Figure 3a). High-content analysis indeed showed that there was a substantial increase in the number of protein inclusions detected per cell for both wild-type and all of the mutants upon hypoglycemia treatment.

### Increased PTEN aggregation in the presence of other aggregated proteins

We and others have shown that aggregation of mutant p53 - the most frequently mutated tumor suppressor in solid cancers – contributes to carcinogenesis by inducing loss-of-function of its tumor-suppressing activity, while acquiring gain-of-oncogenic-function by co-aggregating with various cellular components (De Smet et al., 2017; Xu et al., 2011). Consequently, PTEN destabilizing mutations may occur on a background of other aggregatied proteins, which in turn might be a proteotoxic trigger for PTEN aggregation (De Smet et al., 2016; Gidalevitz et al., 2009).

We first studied the generic effect of an aggregated background on the aggregation behavior of PTEN. To this end, we first created an amyloid-like aggregation background using co-expression of an extended repeat of glutamines (Q97), which forms large amyloid-like fibrillar aggregates. As a control we used a short glutamine repeat (Q20) which remains soluble and does not aggregate. Co-expression of PTEN and Q20 caused a marginal decrease in solubility for all PTEN variants (not shown). Remarkably, upon co-expression with the aggregating Q97 repat, we could detect a significant decrease in solubility for the PTEN R173P mutant, but not for any of the other PTEN variants tested (Supplemental Figure 3a). As both Q20 and Q97 were tagged with mCherry, the aggregation state of both polyQs could be monitored using fluorescent microscopy. This confirmed that while Q20 always retained a diffuse cellular distribution, Q97 localized to large cellular inclusions, indicating that this indeed formed aggregates (not shown).

We next assessed the behavior of PTEN in the presence of the aggregating p53 mutant R175H, which is highly aggregation prone yet forming predominantly soluble oligomeric aggregates in regular cell culture conditions (De Smet et al., 2016). As observed for the polyQs, co-expression of PTEN variants with wild-type p53 as a control only induced a minor decrease in solubility for most PTEN variants (Figure 4c). However, on a background of p53 R175H, however, we again observed that PTEN R173P was the only mutant that showed a significant decrease in solubility. To evaluate whether there was a direct interaction between PTEN R173P and p53 R175H, we performed immunofluorescent staining for both proteins and assessed whether there was co-localization. While staining for p53 R175H generally appeared to be diffuse nuclear or nuclear and cytoplasmic, staining for PTEN R173P was often observed to be localized to inclusions (Figure 4c). Moreover, these inclusions appeared to be devoid of p53, indicating that although the presence of aggregated p53 induces aggregation of PTEN R173P, this effect seemed to be indirect through disturbance of the cellular proteostasis and thus not a result of a direct interaction between aggregating PTEN and p53. The latter was furthermore confirmed by the absence of p53 R175H from the immunoprecipitated fractions of PTEN WT and R173P after co-expression (Supplemental Figure 3c).

### Insoluble PTEN inclusions occur in PDTX models of uterine cancer

Several studies have implicated high incidence of alterations of the *PTEN* gene in uterine and ovarian cancers. Indeed, the incidence of *PTEN* mutations in women diagnosed with uterine and, more specifically, endometrial hyperplasia is one of the highest among analyzed tumors and the most commonly mutated gene identified in uterine cancer (Cancer Genome Atlas Research et al., 2013; Gibson et al., 2016). Based on the high frequency of PTEN inactivating events in uterine cancers, we decided to focus on this cancer type to study the prevalence of PTEN aggregation. We obtained a set of formalin-fixed paraffin-embedded (FFPE) sections from early generation (F2) patient-derived tumor xenograft (PDTX) models of uterine cancer (N=30; Table 2). As protein aggregation usually results in the formation of insoluble intracellular complexes and inclusion bodies, we screened our set of uterine PDTX samples for the presence of PTEN-positive puncta using fluorescent immunohistochemistry (IHC.). This analysis revealed that 26,7% (8/30) of the uterine PDTX samples contained PTEN-positive puncta - most of which in the cytoplasm – usually within the diffuse staining pattern (Figure 5a and Table 2). Sizes of these puncta ranged from 100nm to 5μm, the latter being suggestive for inclusion bodies. Surprisingly, subsequent genotyping of these samples showed that PTEN inclusions were exclusively found in samples that are wildtype for PTEN (Table 2).

**Figure 5.**
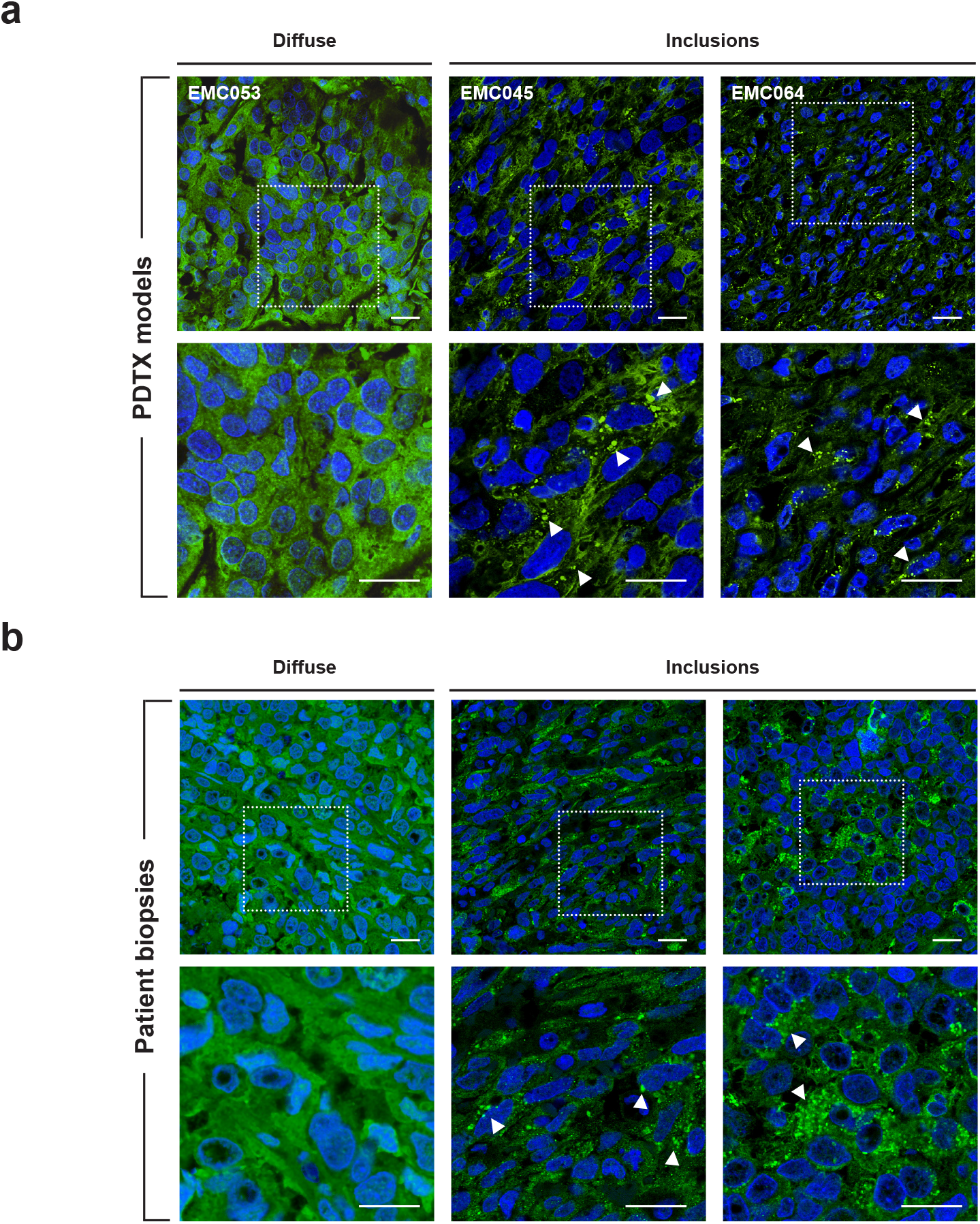
PTEN inclusions in PDTX models and patient biopsies of uterine cancer. Immunofluorescent staining of PTEN in PDTX samples (**a**) and patient biopsies (**b**) of uterine cancer. Confocal overlay images are shown of FFPE sections stained for PTEN (green) and the nuclear dye DAPI (blue) - bottom panels are higher magnification images of the corresponding top panels. Representative images are shown for samples showing diffuse PTEN staining and samples showing PTEN-positive puncta and inclusions (white arrowheads). Scale bars correspond to 20 μm.

**Table 2.** Overview of uterine PDTX sample parameters.

| Tumor ID | Histological subtype | Primary / metastasis / recurrence | PTEN expression<sup>1</sup> | % of cells with PTEN-positive inclusions<sup>2</sup> | Genotype |
| --- | --- | --- | --- | --- | --- |
| EMC011 | Carcinosarcoma, grade 3 | Primary | +/- | ND | WT |
| EMC054 | Carcinosarcoma, grade 3 | Primary | +/- | 25-50% | WT |
| EMC015 | Endometrioid with squamous differentiation, grade 1 | Primary | +/- | <25% | WT |
| EMC016 | Endometrioid with squamous differentiation, grade 2 | Primary | - | ND | A121fs |
| EMC003 | Endometrioid with squamous differentiation, grade 3 | Metastasis/recurrence | - | ND | R130G |
| EMC028 | Endometrioid, grade 1 | Primary | + | ND | WT |
| EMC032 | Endometrioid, grade 1 | Primary | + | ND | F341V |
| EMC001 | Endometrioid, grade 2 | Primary | + | ND | R130L |
| EMC032 | Endometrioid, grade 2 | Primary | + | ND | R130Q |
| EMC034 | Endometrioid, grade 2 | Metastasis/recurrence | - | ND | C71Y |
| EMC046 | Endometrioid, grade 3 | Primary | - | ND | WT |
| EMC068 | Leiomyosarcoma - epithelioid, grade 3 | Metastasis | - | ND | WT |
| EMC031 | Leiomyosarcoma, grade 3 | Metastasis/recurrence | +/- | ND | WT |
| EMC029 | Leiomyosarcoma, grade 3 | Metastasis/recurrence | ++ | 25-50% | WT |
| EMC048 | Leiomyosarcoma, grade 3 | Metastasis/recurrence | ++ | ND | WT |
| EMC059 | Leiomyosarcoma, grade 3 | Metastasis/recurrence | + | ND | WT |
| EMC045 | Leiomyosarcoma, grade 3 | Metastasis/recurrence | + | >50% | WT |
| EMC036 | Leiomyosarcoma, grade 3 | Primary | ++ | <25% | ? |
| EMC043 | Leiomyosarcoma, grade 3 | Primary | ++ | ND | WT |
| EMC041 | Leiomyosarcoma, grade 3 | Metastasis | ++ | <25% | WT |
| EMC050 | Leiomyosarcoma, grade 3 | Metastasis | ++ | ND | WT |
| EMC064 | Leiomyosarcoma, grade 3 | Metastasis | + | >50% | WT |
| EMC007 | Mesonephric, grade 3 | Metastasis/recurrence | +/- | <25% | WT |
| EMC022 | Mesonephric, grade 3 | Primary | - | ND | WT |
| EMC024 | Mesonephric, grade 3 | Metastasis | - | ND | WT |
| EMC042 | Mixed Endometrioid/Clear cell, grade 3 | Primary | +/- | ND | WT |
| EMC002 | Serous, grade 3 | Primary | - | ND | WT |
| EMC008 | Serous, grade 3 | Primary | +/- | ND | R130G |
| EMC053 | Serous, grade 3 | Primary | ++ | ND | WT |
| EMC071 | Undifferentiated uterine sarcoma, grade 3 | Primary | ++ | ND | WT |
| EMC047 | Undifferentiated, grade 3 | Metastasis/recurrence | - | ND | C211fs |

1 – PTEN expression was estimated based on immunofluorescence staining intensity. ++: strong expression; +: weak expression; +/-: mixed expression (partly negative, partly positive); -: no expression.
2 – Percentage of cells showing PTEN-positive inclusions was estimated based on staining of one FFPE section. Given the heterogeneous nature of these phenotypes, these estimations do not necessarily reflect the whole tumor.

To exclude that the occurrence of these inclusions was an artefact e.g. generated by establishing the PDTX models, we obtained FFPE sections from a subset of the original patient biopsies used to generate these PDTX models and stained them for PTEN. Analysis of these samples showed that PTEN expression and subcellular localization (including presence of inclusions) was fully preserved between the primary patient and PDTX samples (Figure 5b).

Poly-ubiquitinated protein aggregates can be selectively recognized by ubiquitin-binding autophagy receptors like p62/SQSTM1 and targeted for degradation by autophagy. Furthermore, p62 was reported to contribute to the formation of inclusion bodies. In line with this, we found that double staining of puncta-containing PDTX samples showed that although the smaller puncta did not show obvious co-staining of p62 and PTEN, we did observe that p62 formed a shell surrounding the larger PTEN inclusion bodies, similar to what was observed for inclusion bodies formed by many aggregating proteins in neurodegenerative diseases (Figure 6a). To further probe the nature of these inclusion bodies, we also stained samples positive for these large PTEN bodies for an aggresome-specific dye (Proteostat (Shen et al., 2011)) and found positive-staining bodies, which is consistent with what we observed for the p62 staining (Supplemental Figure 4).

**Figure 6.**
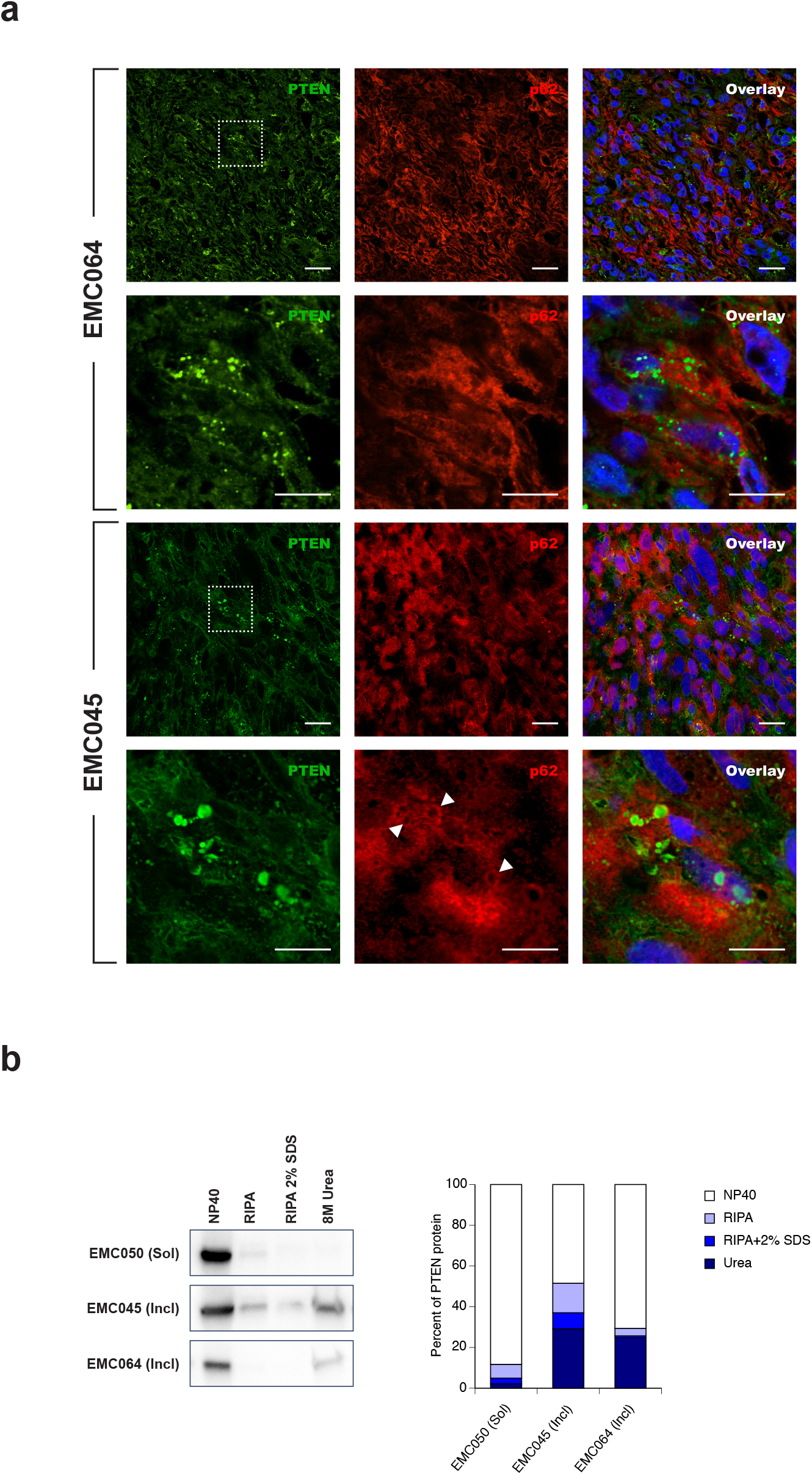
Characterization of PTEN-positive inclusions in PDTX models of uterine cancer. **a.** Immunofluorescent staining of PTEN (green) and p62 (red) in PDTX samples showing PTEN-positive puncta and inclusions. Representative confocal images of the separate PTEN and p62 channels and overlay images with DAPI (blue) are shown for one PDTX sample containing mostly smaller aggregates (EMC064) and one containing also larger, inclusion body-like structures. While no detectable association of the p62 staining was observed for the smaller aggregates, p62 was detected to form a shell around most of the inclusion body-like structures (white arrowheads). For each sample higher magnification images of the boxed areas are shown in the bottom panels. Scale bars correspond to 20 μm and 10 μm in the low and high magnification panels, respectively. **b.** Assessment of PTEN protein solubility using the serial extraction assay in a selected set of PDTX samples. While for the sample showing diffuse PTEN staining (EMC050) most of the PTEN protein was extracted with the mildest extraction buffer (NP40), a considerable portion of the protein was only extracted with the most denaturing buffer (8M urea) for the inclusion-containing samples (EMC045 and EMC064). Left panels show the Western blots of the fractionated samples while the right panel shows the densitometric quantification of these blots as percentage of PTEN protein per solubility bin.

Finally, to assess the aggregation status biochemically, we performed the serial solubility protein extraction assay, as described above. Upon serial extraction of a PDTX sample showing a diffuse immunofluorescent staining pattern, most of the PTEN protein (>85%) was detected in the most soluble NP40 fractions, while only background amounts (around 2%) were detected in the most insoluble urea fractions (Figure 6b). Conversely, similar analysis of two samples showing PTEN-positive inclusions - the 2 models in which >50% of cells show inclusions (Table 2) were selected to minimize the dilution effect - showed a marked shift in PTEN solubility, mainly towards the most insoluble urea fractions. Indeed, extraction of inclusion-containing samples generated urea fractions containing >20% of the total PTEN protein content, showing that the presence of PTEN-positive inclusions correlates with a shift towards insoluble, aggregated PTEN protein (Figure 6b).

### Exploratory retrospective study of PTEN aggregation in endometrial cancer samples

Finally, we wanted to investigate the potential relevance of PTEN aggregation for the clinical outcome in endometrial cancer patients. To this end, we obtained FFPE material from a small exploratory cohort of 20 cases of type I, and 65 cases of type II endometrial cancer (Table 3). Type I endometrial carcinoma are generally slow growing and well differentiated (grade I, II), whereas type II cancers are more aggressive and likely to spread and undifferentiated (grade III). On a molecular level, PTEN inactivation is observed in about 35-50% of type I cases, while type II cases show very few genetic alterations with PTEN inactivation being present in only around 10% of cases (Lax, 2004).

**Table 3.** Patient characteristics overview

|  |  |
| --- | --- |
| Endometrial cancer Patients | 85 (100%) |
| Age at diagnosis (years) | 66 mean (36 – 89) |
| Progression free survival (months) |  |
| Type I | 72.3 mean (1 – 119) |
| Type II | 76.5 mean (0 – 278) |
| Overall survival (months) |  |
| Type I | 77.7 mean (1 – 119) |
| Type II | 82.17 mean (8 – 278) |
| FIGO stage |  |
| Type I |  |
| FIGO I | 14 |
| FIGO II | 2 |
| FIGO III | 4 |
| Type II |  |
| FIGO I | 27 |
| FIGO II | 7 |
| FIGO III | 12 |
| FIGO IV | 15 |
| unknown | 4 |
| Histological Grading |  |
| Type I G1 | 12 |
| Type I G2 | 8 |
| Type IIG2-3 | 2 |
| Type IIG3 | 60 |
| Type IIMV | 3 |
| Histo Type |  |
| Type I endometrioid | 20 |
| Type IIendometrioid | 20 |
| Type IIendometrioid, serous | 13 |
| Type IIendometrioid, clear cell | 1 |
| Type IIserous | 17 |
| Type IIclear cell | 8 |
| Type IIcarcinosarcoma | 4 |
| Type IImucineus | 2 |

To quantify the level of PTEN aggregation in these samples, we employed the so-called Seprion-ELISA, which was previously shown to yield highly reliable quantification of amyloid like aggregation in tissues (Maritschnegg et al., 2018). Similar to what was observed *in vitro* and in PDTX models, only 3/20 (15%) of type I tumors showed an absorbance above the threshold in the Seprion-ELISA, whereas background levels were measured in 17/20 (85%). In contrast, in type II cases 34/65 (52%) scored positive. In 13/34 (38.2%) of these cases, the detected levels were beyond an absorbance of 1.0. The median absorbance measured in the positive type I cases was 0.269 (range: 0.267 - 0.531), compared to a median absorbance of 0.649 (range: 0.251 - 3.692) in the type II cases (Figure 7a). The *PTEN* mutation status was known in 58 of the type II cases. The presence of a mutation was confirmed in 8/58 (13.8%) of them. However, in this subset of patients, no correlation was observed between the presence of a *PTEN* mutation status and the Seprion aggregation load (Fisher’s exact test two-tailed P value equals 0.7073). Furthermore, the *TP53* mutation status was also known in 58 cases. No correlation between the presence of a *TP53* mutation and PTEN aggregates was observed (Fisher’s exact test two-tailed P value equals 1.0).

**Figure 7.**
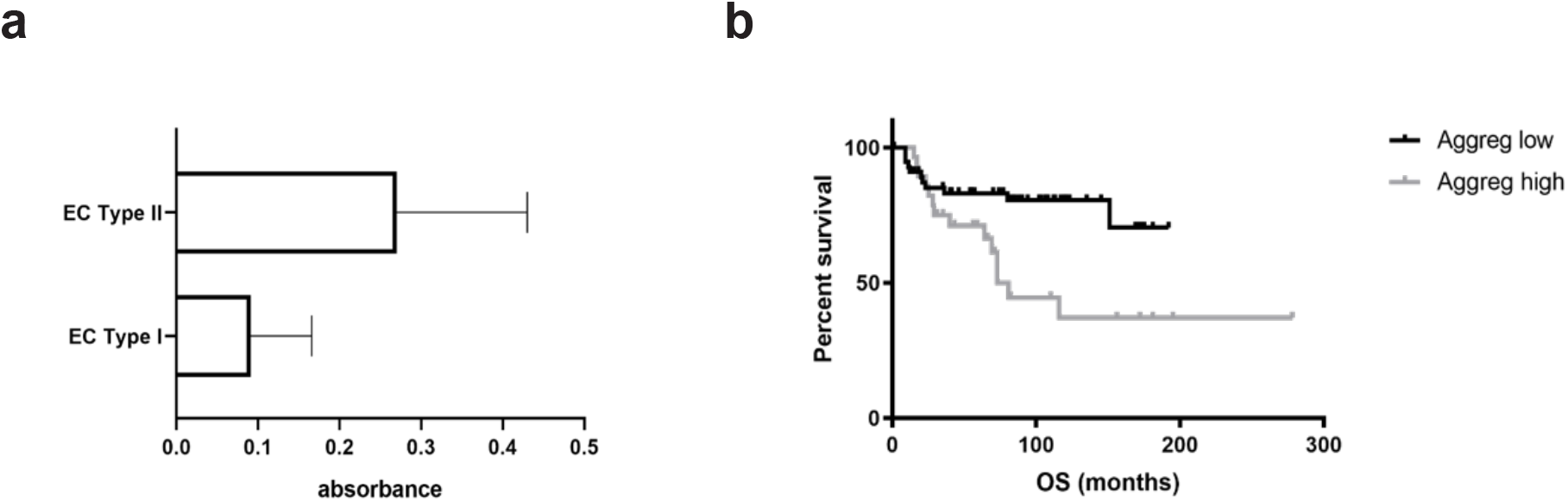
Exploratory retrospective study of PTEN aggregation in endometrial cancer samples. **a.** Comparison of the levels of aggregated PTEN between type I and type II EC measured in the Seprion-ELISA. The measured absorbance was significantly higher in type II cancers (T-test two tailed P=0.0106, median absorbance with 95% CI). **b.** Survival curve for all type I and type II EC patients included in this study, subdivided into a group with high and low PTEN aggregation load, as determined by Seprion-ELISA. Patients with PTEN aggregation low tumors have a significantly improved OS (log rank p=.0103), compared to those with a high aggregation load.

Finaly, the Kruskall-Wallis test was used to compare the PTEN aggregation level between cancer stages, determined according to the International Federation of Gynecology and Obstetrics (FIGO) criteria. No significant correlation between the PTEN aggregation load and FIGO-stage was observed (P=0.6856). However, when we stratified the patients by PTEN aggregation as quantified by the Seprion-ELISA assay, we observed a significant decrease in the overall survival rate of the patients that carry PTEN aggregates compared to those that do not (Mantel-Cox, P=0,0103, Hazard ratio 0,3307; Figure 7b).

## Discussion

We find that wild-type PTEN aggregates into amyloid-like inclusions in endometrial tumors, an observation which may correlate to a decrease in patient survival. These data suggest that aggregation of the tumor suppressor PTEN constitutes a novel and distinct structural mechanism driving oncogenesis independently of genetic tumor suppressor mutation. Aggregation of tumor suppressors including p53 and axin has previously been demonstrated and for both proteins aggregation-associated oncogenic gain-of-function has been reported (Ano Bom et al., 2012; Anvarian et al., 2016; De Smet et al., 2017; Ghosh et al., 2017; Xu et al., 2011). However, in these contexts aggregation of the tumor suppressors is generally considered to be a consequence of mutation: cancer mutations affect the native structure of tumor suppressors causing protein misfolding which in turn favors aggregation.

Mechanistically, amyloid-like aggregation is driven by short aggregation prone segments (APRs) within protein sequences (Rousseau et al., 2006). When identical APRs from different protein chains interact, they are stabilized by ß-strand-interactions, which can grow until they form large insoluble ß-sheet structured assemblies. In native proteins these APRs are generally buried inside the hydrophobic core of the protein and therefore protected from aggregation. Cancer mutations that destabilize the native structure therefore favor aggregation by increasing solvent exposure of these APRs. However, these same APRs that trigger aggregation in mutant tumor suppressors are also present in the wild-type protein suggesting that under conditions of increased physiological stress wild-type proteins are also likely to aggregate. In fact, wild-type protein aggregation is the most prevalent scenario in the most common amyloid-associated neurodegenerative diseases, such as Alzheimer or Parkinson disease whereas familial forms of these diseases are much less common (Bertram and Tanzi, 2005). The main risk factor for protein aggregation in neurodegenerative diseases is ageing (Rhodes and Ritz, 2008). The most probable dominant cause triggering the onset of protein aggregation in this context is the decline in proteostatic control in ageing cells, including reduced Hsf-1 dependent chaperone upregulation, proteasomal regulation or autophagy (Douglas and Dillin, 2010; Kennedy et al., 2014; Lopez-Otin et al., 2013). This results in an increased risk of protein aggregation by a combination of increased frequency of stochastic misfolding events and a decreased ability to handle these. The origin of proteostatic decline in turn is thought to result from accumulated oxidative damage impairing fundamental mechanisms such as mitochondrial function thereby gradually restricting resources to revert the entropic degeneracy of cellular integrity (Hohn et al., 2017).

Here we show that aggressive type II endometrial tumors may evade wild-type PTEN function by allowing for its aggregation. In a small exploratory cohort of type I and II endometrial cancer patients, we assessed the correlation between PTEN aggregation and clinical outcome. PTEN aggregation was predominately observed in type II tumors, where it appeared to be correlated with reduced overall survival. Given the small sample size and the exploratory nature of the cohort, the weight of these findings can be questioned, but the observation certainly points towards the need for a more thorough investigation. Similar findings were furthermore made in patient-derived murine xenograft samples of endometrial cancers and their original human tumour counterparts. The latter could be an important observation as it suggests that the phenotype is driven, at least in part, by the intrinsic nature of the tumor cell and not only by its environment.

*In vitro* characterization showed that PTEN wild-type aggregation can be induced in cancer cells by both intrinsic proteotoxic stress (i.e. by proteasomal inhibition or coexpression of other aggregating proteins) as well as by applying micro-environmental stresses encountered during tumor growth such as hypoxia and hypoglycemia. Indeed, cancer cells growing in the tumor environment are exposed to several stresses affecting proteostatic fitness. First, cancer cells gradually accumulate both driver and passenger mutations in various proteins resulting in an increased and continuous production of misfolded proteins. While cancer cells benefit from mutations in tumor suppressors they also have to avoid dying from massive protein misfolding. As a result, cancer cells adapt Hsf-1 to the extent that many tumor cells over time develop chaperone addiction as is apparent from their inability to survive Hsf-1 inhibition (Dai et al., 2007; Mendillo et al., 2012; Whitesell and Lindquist, 2005). In addition to this internal proteostatic stress, tumor growth also results in micro-environmental stress including hypoxia and hypoglycemia. This situation of chronic proteostatic stress in tumors suggests that our observation of wild-type PTEN aggregation in human cancer cells might not be a unique case but that proteostatic stress in cancer cells might - beyond mutation-be a generic mechanism favoring the inactivation of other wild-type tumor suppressors. Indeed, similar observations were made for the wild-type p53 protein, for which inclusions were found in a range of human tumors (De Smet et al., 2017), supporting this hypothesis.

Destabilizing and truncating cancer mutations in PTEN, finally, significantly aggravated the aggregation propensity of the protein *in vitro* as compared to wild-type. Under stressed conditions, these mutants undergo severe aggregation as evidenced by the formation of large inclusions that show many features of amyloid aggregates. However, amyloidogenesis has been shown to act tumor suppressive and tumor cells have been shown to rewire their signaling pathways pro-actively to avoid formation of amyloid(-like) aggregates (Dai 2015). In line with this, we found no evidence of PTEN aggregation in PTEN mutant patient-derived samples, although evaluation of larger cohorts is required to further substantiate these observations.

## Materials and methods

### In silico analysis of aggregation, stability and structure of PTEN

The aggregation propensity of Osgin-1, PTEN WT and its mutants was analysed with TANGO (Fernandez-Escamilla et al., 2004), an algorithm to predict aggregation-nucleating sequences in proteins. The effect of the mutations on PTEN stability was analysed by calculating the change in free energy (ΔΔG) upon mutation with the FoldX force field (Schymkowitz et al., 2005). Structural changes of PTEN due to mutations were inspected with YASARA (Krieger et al., 2002).

### Recombinant expression, biophysical characterization and transmission electron microscopy of N-PTEN

Plasmid 821 pGEX2T PTEN 1-274 (N-PTEN) was a gift from William Sellers (Addgene plasmid # 10741) (Ramaswamy et al., 1999). The GST tag in this construct was replaced by a 6xHis tag using the Q5^®^ Site-Directed Mutagenesis Kit (New England Biolabs). N-PTEN was recombinantly expressed in E. coli and purified from inclusion bodies using a HisTrap column (GE Healtcare Life Sciences) in 8M urea.

Aggregation of N-PTEN was assessed using the amyloid-specific dye Thioflavin T (ThT) and the hydrophobic dye 1-anilino-8-naphthalene sulfonate (ANS). 40 μl protein samples (50 μM) were placed into a flat-bottom 384-well microclear plate (Greiner, Frickenhausen, Germany) and the dyes were added to a final concentration of 10 μM. A FLUOstar OPTIMA multidetection microplate reader (BMG labtech, Offenburg, Germany) was used to measure fluorescence.

For TEM imaging, Formvar film coated 400-mesh copper grids (Agar Scientific Ltd., England) were first glow-discharged to improve adsorption efficiency. Next, 10 μL of each sample was adsorbed for 10 min and afterwards the grids were washed by contact with 3 drops of ultrapure water. Finally, a negative staining was performed by contact with one drop of uranyl acetate (2% w/v) for 1 min. The grids were examined using a JEM-1400 transmission electron microscope (Jeol, Japan) at 80 keV.

### Immunofluorescence on PDTX samples and patient biopsies

FFPE sections from PDTX samples (N=33) and patient biopsies (N=8) were collected and provided by the TRACE platform, KU Leuven - UZ Leuven (www.uzleuven-kuleuven.be/lki/trace). Sections were stained with the following antibodies: mouse monoclonal anti-PTEN (Dako), rabbit polyclonal anti-p62 (Enzo Life Sciences) and rabbit polyclonal anti-Osgin-1 (Bioss Antibodies). Deparaffinization and heat-induced epitope retrieval was performed using the Target Retrieval Solution (pH 9.0; Dako) at 95°C for 40mins. For LCP dye staining, slides were deparaffinised and incubated with 3 μM LCP in PBS for 30 mins at RT. Next, slides were washed in PBS and mounted with Dako fluorescence mounting medium.

### Cell culture and transient transfection

HeLa and PC-3 cells were cultured in Dulbecco’s modified eagle medium (DMEM) and Ham’s F-12K (Kaighn’s) medium (both from Life Technologies), respectively, supplemented with 10% FCS, non-essential amino acids and sodium pyruvate, at 37°C in 5% CO2. For transient transfections in 6-well culture plates, 4×10^5^ HeLa or 6×10^5^ PC-3 cells were plated per well. The following day, 1,5 μg plasmid DNA was transfected using the Lipofectamine 2000 transfection reagent (Life Technologies) according to manufacturer’s protocol. For transient transfection in 96-well culture plates, 10^4^ HeLa cells were plated per well. Treatment of cells with the proteasomal inhibitor MG132 was performed for 16 hrs starting 6 hrs after transfection. Hypoglycemia and NiCl_2_ treatments were performed for 40 hrs.

### Immunofluorescence

HeLa or PC-3 cells were plated (3×10^4^ and 4×10^4^, respectively) and transfected on glass 8-well chamber slides. 24 hrs after transfection, the cells were fixed with 4% formaldehyde (5 min, RT), washed in Tris Buffer Saline pH 7.4 (TBS), permeabilized in 0.2% Triton X-100 in TBS (5 min, RT) and blocked (2 x 15 mins, RT) in a blocking buffer (5% BSA in TBS). The primary antibodies, mouse anti-HA tag (Biolegend; 1/750), rabbit anti-Flag tag (Cell Signalling; 1/1000), were diluted in blocking buffer and incubated 1 h at RT. The secondary antibodies (goat anti-mouse Alexa Fluor 594 and goat anti-rabbit Alexa Fluor 488) were diluted 1:1000 and incubated for 1 h at RT. The slides were mounted with ProLong Gold Antifade Reagent containing DAPI (Invitrogen). Images were acquired using a Leica SP8x confocal microscope. LCP staining was performed by adding the dye 1/1000 after the immunofluorescence procedure.

### High content analysis of the PTEN aggregating mutants

HeLa cells were seeded and transfected in 96-well plates as described above. 24 h after transfection, cells were washed in TBS, fixed in 4% formaldehyde and stained for immunofluorescence as described above. Nuclei were stained with DAPI diluted 1:10.000 and incubated together with the secondary antibody. To quantify the cells with protein inclusions, plates were analysed using the IN Cell analyser 2000 (GE Healthcare) high-content analysis system. Image acquisition was done using a 60x objective. For image analysis, a custom algorithm was developed on the IN Cell Developer Toolbox (GE Healthcare) platform.

### Serial protein extraction

At indicated time-points after transfection, HeLa cells were lysed in NP40 buffer (150mM NaCl, 50mM Tris HCl pH8, 1% IGEPAL(NP40), 1x Complete inhibitor (Roche), 1U/μl Universal Nuclease (Pierce)). Extracts were kept on ice for 45 mins after which the insoluble fraction was collected by centrifugation for 10 mins at 13,3×10^3^ rpm at 4°C. The supernatant was collected as the NP40 soluble fraction, while the pellet was redissolved in RIPA buffer (Pierce). Redissolved extracts were again incubated on ice for 45 mins followed by centrifugation as described before. Pellets were next redissolved in RIPA buffer containing 2% SDS and after an incubation and centrifugation step, the remaining pellets were dissolved in 8M Urea. The serial extraction thus generates four protein fractions with different solubilities: the NP40, RIPA, RIPA+2%SDS and Urea soluble fractions. Protein extracts were fractionated by SDS-PAGE and subsequently transferred by electroblotting (fixed current 0.4 A) on a nitrocellulose membrane (BioRad). The membrane was incubated in 5% dried non-fat milk powder dissolved in TBS-Tween 0,1% (TBST) for 1 h at room temperature (RT) and subsequently incubated with primary anti-HA antibody (Biolegend) followed by incubation by secondary goat HRP-conjugated anti-mouse IgG (Promega). Proteins were visualized using chemiluminescence detection reagent (ECL, Millipore). Protein levels in each solubility fraction were quantified densitometrically and expressed as percentage of the total level (i.e. sum of the four fractions).

### Filter trapping

Transfected cells were lysed at indicated time-points in a RIPA buffer containing 1% SDS. Whole, non-cleared lysates were next filtered through Spin-X 0,22μm cellulose acetate filters by centrifugation at 13,3×10^3^ rpm for 15 mins. Proteins trapped on the filter were extracted by a serial approach: (i) first, filters were incubated in RIPA buffer containing 2% SDS at 95°C for 10 mins, followed by a centrifugation step as before and (ii) next, the filters were incubated in 8M Urea for 30 mins at RT and centrifuged again. Different fractions were fractionated and quantified as described above.

### Immunoprecipitations

For co-immunoprecipitations, PC-3 cells were (co-)transfected in 6-well plates. 24 hrs after transfection, cells were lysed in RIPA buffer supplemented with 1x Complete inhibitor (Roche) and 1U/μl Universal Nuclease (Pierce). Lysates were incubated with a rabbit anti-HA antibody (1/100, Cell Signalling) overnight at 4°C on a rotor. In parallel, magnetic beads (Dynabeads, Invitrogen) were blocked in 5% BSA/TBS. Next day, the antibodies in lysates were trapped on the magnetic beads, and washed extensively in RIPA buffer. Bound proteins were dissociated from the antibodies by competitive elution with HA peptide (0,5 mg/mL) and analysed by Western blotting.

For immunoprecipitations with the OC antibody, a similar approach was followed with the following adaptations: cells were lysed in NP40 buffer (see above) and immunoprecipitated proteins were eluted by boiling the magnetic beads in 1x Western blot sample buffer.

### Patient samples

Fresh frozen tumor tissue specimens of 85 women with endometrial cancer, which have been collected for diagnostic purposes at the University Hospitals Leuven were used in the present study. All samples were collected and analyzed in accordance with the institutional review board (S61589). Informed consent was obtained from each participant. Of those tissues, 20 were classified as type I endometrial cancer based on grade and histology (grade I & II, endometrioid), and the remaining 65 were classified as type II endometrial cancer (grade III, endometrioid, serous, clear cell, undifferentiated). Molecular information as well as molecular classification of the tissues was available from the publication “Amplification of 1q32.1 refines the molecular classification of endometrial carcinoma” (Depreeuw et al., 2017). A tumor cell content of at least 50% was ensured by a pathologist via cryo-sectioning and subsequent H&E staining. Furthermore, the tissue was evaluated for signs of necrosis.

### Seprion-ELISA

The tumor tissue was weighed and ice-cold NP-40 buffer including a protease inhibitor cocktail (cOmplete inhibitor, Roche) was added in order to prepare a 2.5% (w/v) lysate. Tissues were homogenized using the FastPrep-24 instrument (MP Biomedicals). After incubation on ice for 30min, the lysates were frozen on dry ice and stored at −80°C.

The PTEN aggregation load was determined using the Seprion-ELISA (Microsens) as described in (Maritschnegg et al., 2018). This test is based on a polyionic, high-molecular-weight ligand, coated to the surface of the ELISA plate. In combination with the provided Seprion capture buffer, only aggregated, and/or amyloid proteins can bind to the ligand, whereas naturally occurring multimers are washed away. Using an anti-PTEN specific primary antibody, the Sandwich ELISA therefore allows detecting solely aggregated PTEN. This test not only has the advantage of high-throughput analysis, but it is also the only available test that allows a quantitative detection of aggregates from cell lysates, which is why this assay was chosen for this particular study. A 1:20 dilution of the lysate was applied to the ELISA and measured in triplicates using a plate reader (ELx808, Beun DeRonde Serlabo BVBA). The signal intensity is proportional to the bound mass of aggregated PTEN. The primary antibody used for the identification for PTEN aggregates was anti-PTEN 26H9 (Cell Signaling Technology), diluted 1:100. Additionally, a negative control was included in every experiment and measured in triplicates. The highest absorbance measured in one of the negative controls, plus three times the standard deviation, was used as the threshold for a positive signal.

## Acknowledgements

The Switch Laboratory was supported by grants from the European Research Council under the European Union’s Horizon 2020 Framework Programme ERC Grant agreement 647458 (MANGO) to JS, the Flanders institute for biotechnology (VIB), the University of Leuven (“Industrieel Onderzoeksfonds”), the Funds for Scientific Research Flanders (FWO), the Flanders Agency for innovation by Science and Technology (IWT, SBO grant 60839) and the Federal Office for Scientific Affairs of Belgium (Belspo), IUAP, grant number P7/16. FC, EM and FDS were supported by fellowships from FWO. FA is a senior researcher of the FWO. The Leica SP8x confocal microscope was provided by InfraMouse (KU Leuven-VIB) through a Hercules type 3 project (ZW09-03). The InCell Analyzer 2000 was made accessible by the Light Microscopy and Imaging Network (LiMoNe), which is part of the VIB Bio-Imaging core at the KULeuven. The authors thank Álvaro Cortés Calabuig from the Genomics Core Leuven, UZ Leuven, Belgium, for his assistance with analyzing sequencing data.

## Author Contributions

Conceptualization, F.C, A.S., J.S., F.R.; Methodology, F.C., G.D.B., A.S.; Software, F.C., G.D.B., J.S., F.R.; Formal Analysis, F.C., G.D.B., J.S., F.R.; Investigation, F.C., A.S., M.S.R., M.R., E.M., F.D.S, J.D.; Resources, E.H., F.A., D.L., K.P.R.N., P.H.; Writing - Original Draft, F.C., J.S., F.R.; Supervision, J.S., F.R.; Project Administration, F.C.; Funding Acquisition, J.S., F.R.

## Supplemental figure legends

**Supplemental Figure 1:**
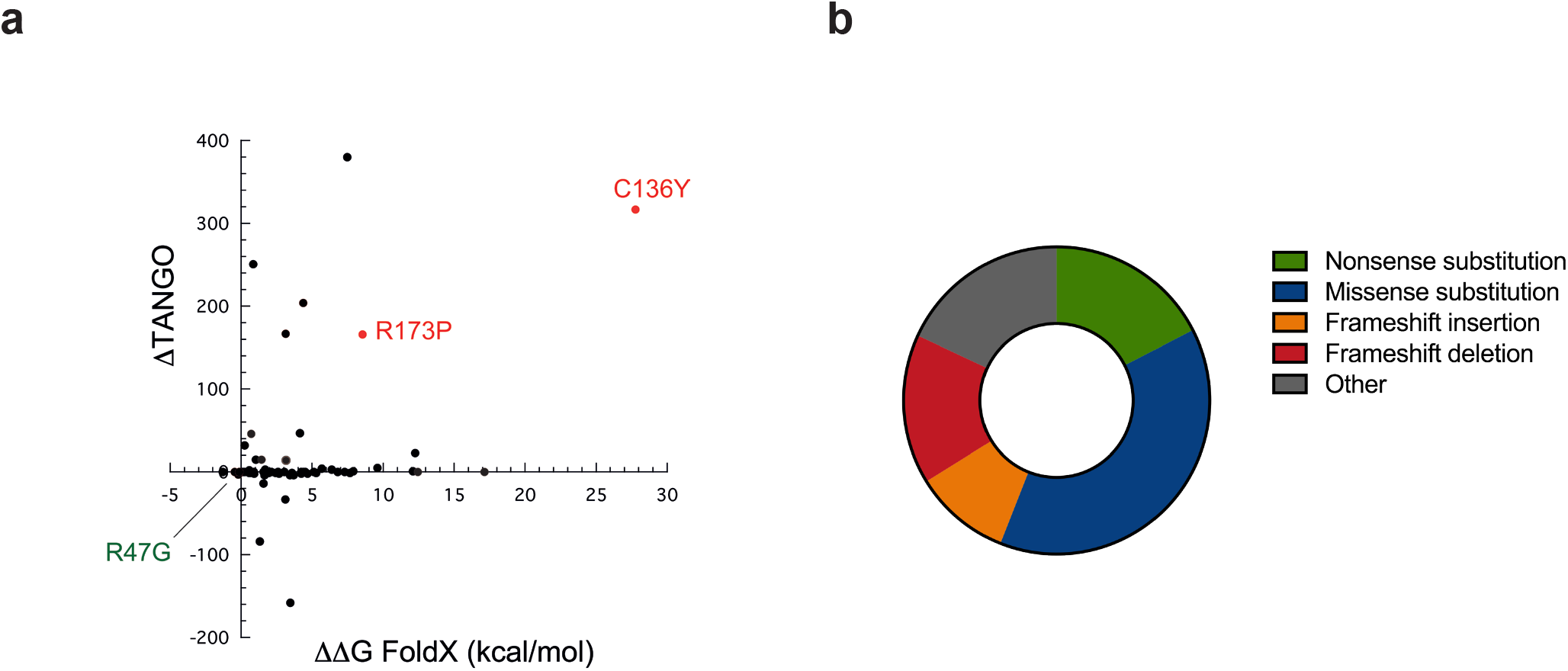
*In silico* analysis of aggregation propensity and stability of disease-associated PTEN mutants. **a.** Mutant aggregation and stability spectrum (MASS) plot for all disease-associated PTEN mutants. The change in the thermodynamic stability (ΔΔG) associated with mutation was plotted versus the corresponding change in intrinsic aggregation propensity (ΔTANGO). This scatter plot shows that the majority of the PTEN mutations do not affect the intrinsic aggregation propensity of the protein, but rather increase aggregation by reducing the thermodynamic stability of the protein. The mutations used in this study are indicated in red (mutants with increased aggregation propensity) and green (control mutant). **b.** Distribution of cancer-associated PTEN mutation types showing that nonsense and frame-shift mutations are at least as predominant as missense mutations.

**Supplemental Figure 2:**
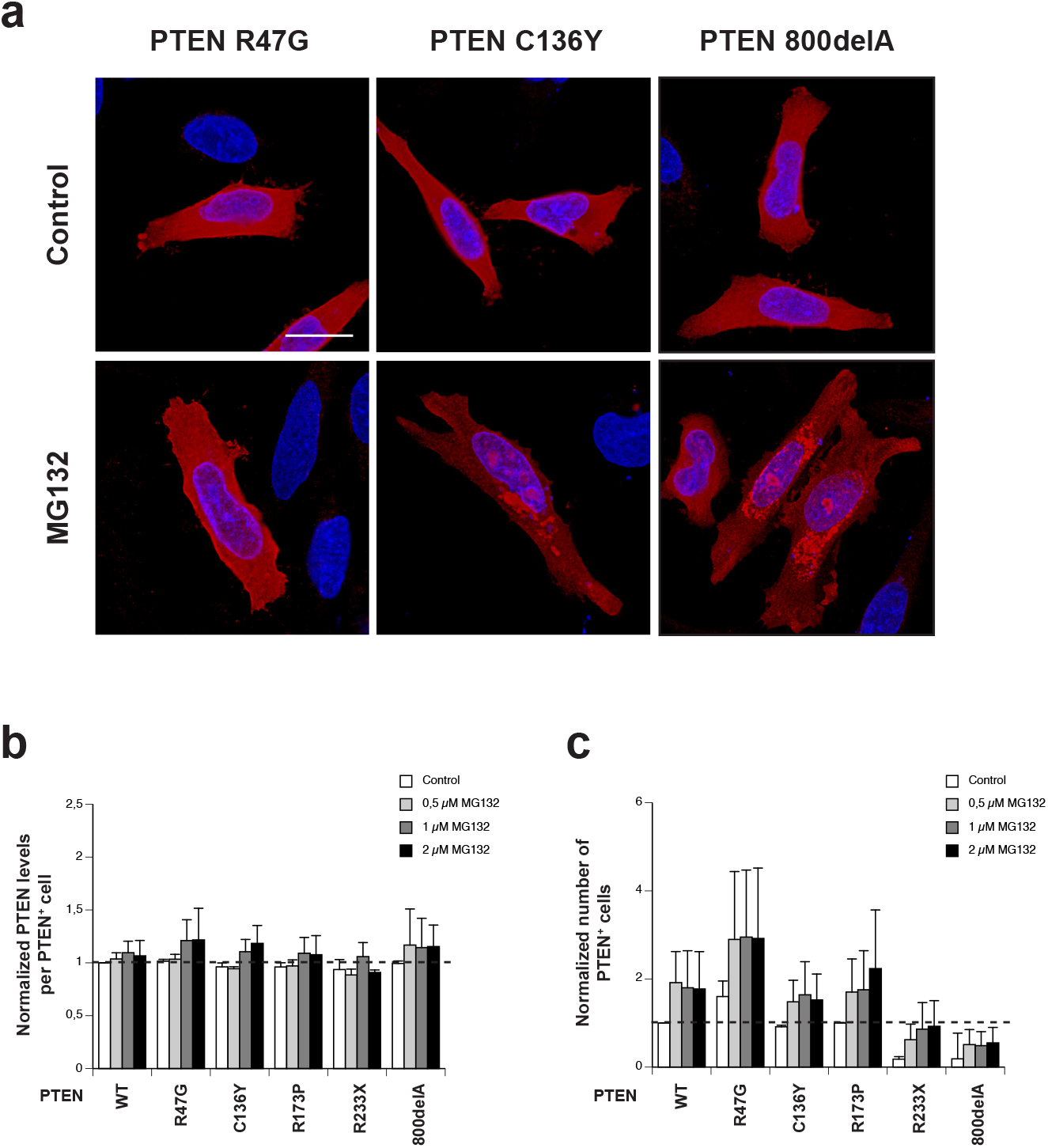
Cellular distribution of PTEN. **a.** Immunofluorescence of the control mutant PTEN R47G, the destabilizing missense mutant C136Y and the frame-shift mutant 800delA in HeLa cells under optimal cell culture conditions (top row) and after proteasomal inhibition with MG132 treatment (bottom row). Under control conditions all PTEN variants shows a diffuse distribution throughout the cell. However, upon MG132 treatment, the staining of the PTEN mutants C136Y and 800delA becomes localized to protein inclusions, while most of the PTEN R47G control mutant protein retains its diffuse staining pattern. Scale bar represents 20 μm. **b.** High-content data on the normalized average levels of PTEN per cell. This analysis did not reveal any differences between the different PTEN variants under any of the conditions tested. **c.** High-content data on the normalized number of PTEN-positive cells showing that under control conditions all full-length PTEN variants are detected in a comparable number of cells. The nonsense and frame-shift mutants, however, are detected in >5 less cells under control conditions. Treatment of transfected cells with MG132 resulted in an increase of PTEN-positive cells for all variants.

**Supplemental Figure 3:**
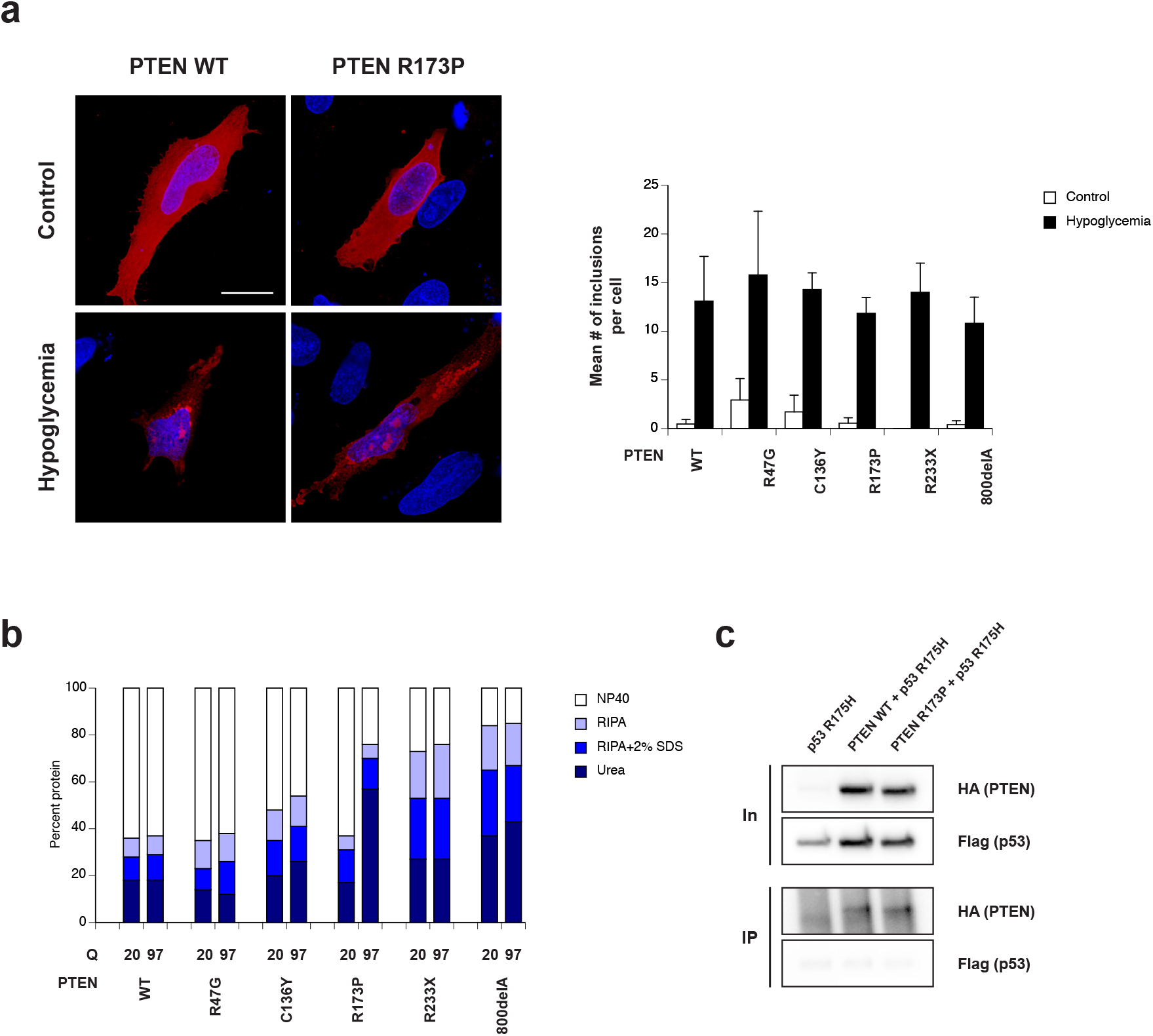
PTEN aggregation upon hypoglycemia treatment or co-expression of other aggregation-prone proteins. **a.** (left panels) Immunofluorescence of PTEN WT and the missense mutant R173P under control conditions (top row) and after treatment with hypoglycemia for 48 hrs (bottom row). Under control conditions both PTEN variants shows a diffuse distribution throughout the cell. However, upon hypoglycemia treatment, the staining of both PTEN WT and R173P becomes localized to protein inclusions. Scale bar represents 20 μm. (right panel) High-content analysis of protein inclusion formation of PTEN WT and the selected mutants in HeLa cells under control conditions and upon treatment with hypoglycemia for 48 hrs. Treatment with hypoglycemia resulted in a large increase in the average number of protein inclusions per cell for all PTEN variants. **b.** Plot of the distribution of PTEN protein over the tested solubility bins upon co-expression with the non-aggregating short polyglutamine stretch Q20 and the aggregating long stretch Q97 in the serial extraction assay. This analysis showed that under these conditions only PTEN R173P showed a clear reduction in protein solubility when co-expressed with Q97. **c.** Co-immunoprecipitation of p53 R175H with PTEN WT or R173P. The immunoprecipitated fractions using PTEN WT and R173P as bait were probed for the presence of p53 R175H. However, no specific p53 signal could be detected in both conditions.

**Supplemental Figure 14.**
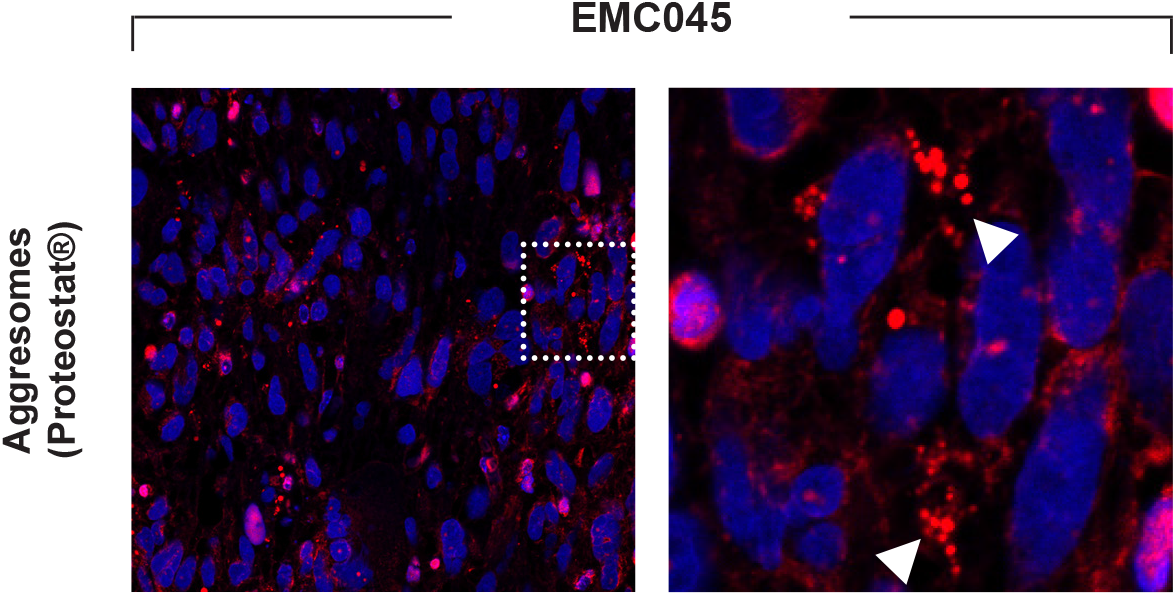
Aggregate-specific marker staining in PTEN inclusion bodypositive PDTX samples. Staining for the aggresome specific-dye Proteostat®, a red fluorescent molecular rotor dye to specifically detect denatured protein cargo within aggresomes and aggresome-like inclusion bodies.

